# HPF1-dependent PARP activation promotes LIG3-XRCC1-mediated backup pathway of Okazaki fragment ligation

**DOI:** 10.1101/2020.10.21.348292

**Authors:** Soichiro Kumamoto, Atsuya Nishiyama, Yoshie Chiba, Ryota Miyashita, Chieko Konishi, Yoshiaki Azuma, Makoto Nakanishi

**Author notes:** To whom correspondence should be addressed., Correspondence may also be addressed to Atsuya Nishiyama.

## Abstract

DNA Ligase 1 (LIG1) is known as the major DNA ligase responsible for Okazaki fragment joining. Recent studies have implicated LIG3 complexed with XRCC1 as an alternative player in Okazaki fragment joining in cases where LIG1 is not functional, although the underlying mechanisms are largely unknown. Here, using a cell-free system derived from *Xenopus* egg extracts, we demonstrated the essential role of PARP1-HPF1 in LIG3-dependent Okazaki fragment joining. We found that Okazaki fragments were eventually ligated even in the absence of LIG1, employing in its place LIG3-XRCC1 which was recruited onto chromatins. Concomitantly, LIG1 deficiency induces ADP-ribosylation of histone H3 in a PARP1-HPF1-dependent manner. The depletion of PARP1 or HPF1 resulted in a failure to recruit LIG3 onto chromatin and a subsequent failure in Okazaki fragment joining in LIG1-depleted extracts. Importantly, Okazaki fragments were not ligated at all when LIG1 and XRCC1 were co-depleted. Our results suggest that a unique form of ADP-ribosylation signalling promotes the recruitment of LIG3 on chromatins and its mediation of Okazaki fragment joining as a backup system for LIG1 perturbation.

## INTRODUCTION

Lagging strand synthesis is a tightly coordinated stepwise reaction involving multiple proteins, including PCNA, which is an essential component of DNA replication (1–3). When DNA polymerase delta (POL δ) reaches the end of a nascent Okazaki fragment, it generates a 5’ flap structure by displacing the 5’ end of the preceding fragment. The displaced 5’ flap is cleaved by nucleases, such as FEN1 or DNA2, and generates a ligatable nick (4–6). Lagging strand synthesis is completed by joining millions of Okazaki fragments through the action of DNA ligases, and a failure of Okazaki fragment joining results in an enormous amount of gap/nick formation on genomic DNA. Therefore, ligation of Okazaki fragments by DNA ligase must be strictly controlled to reliably maintain genome stability.

Vertebrates have three different types of DNA ligases, LIG1, LIG3, and LIG4 (7). While LIG1 is the major ligase functioning in DNA replication, LIG4 plays an essential role in non-homologous end-joining (NHEJ), the pathway for DNA double-strand break repair (DSBR). In budding yeast, the CDC9 gene, a homolog of human LIG1, is essential for viability (8, 9). Cdc9 physically interacts with PCNA via its conserved PCNA interacting peptide (PIP) box motif at the N-terminus and localizes at sites of DNA replication, catalyzing the ligation of Okazaki fragments (10–12). In vertebrates, LIG1 also localizes at DNA replication foci and functions as the main DNA ligase responsible for Okazaki fragment joining (10, 13). Unlike yeast, however, LIG1 is not essential for cell survival (14–17), implying the existence of a compensatory pathway other than that of LIG1.

LIG3 is a DNA ligase characteristic of vertebrates. Nuclear LIG3 forms a tight complex with its partner protein X-ray repair cross-complementing protein 1 (XRCC1) and is involved in nucleotide excision repair (NER), single-strand break repair (SSBR), and base excision repair (BER) (7, 18–20). It has been also suggested that LIG3 might be responsible for alternative Okazaki fragment ligation in the absence of LIG1. Systematic studies using a chicken B cell line demonstrated that even LIG1/LIG4 double knockout cells are viable (15), suggesting that LIG3 may function in DNA replication in the absence of LIG1. The question arises as to how LIG3 is specifically localized to DNA replication sites in the absence of LIG1 during S phase progression, because the domain structures of LIG1 and LIG3 are very different, especially given that LIG3 lacks a PIP-box. Intriguingly, XRCC1 has been shown to associate and co-localize with PCNA during the S phase (21, 22), but other studies have concluded that poly (ADP-ribose) (PAR) synthesis is important for LIG3-XRCC1 recruitment at chromosomal DNA break sites (23–25).

Mono- and poly-ADP-ribosylation are catalyzed by poly (ADP-ribose) polymerase (PARP) family proteins (26), and degraded by terminal ADP-ribose glycohydrolases (TARG) and PAR glycohydolases (PARG) (27, 28). Among the PARP family, PARP1/2 plays a critical role in the SSBR, and DSBR, the regulation of the stability of DNA replication forks, and maintenance of chromatin structures (29, 30). Upon DNA damage, PARP1 rapidly recognizes and binds to a nicked or a gapped single-stranded DNA break via its N-terminal zinc finger domains (31). DNA binding of PARP1 allosterically activates it and promotes its autoPARylation (32), which acts as a platform for tethering SSBR machinery (30). Various PARP1-interacting regulatory factors also play an important role in the regulation of PARP1 activity (30). In response to DNA damage, histone PARylation factor 1 (HPF1) directly binds to the catalytic domain of PARP1/2 via its C-terminal domain (33–35). After the interaction, HPF1 and PARP1/2 jointly form an active site, leading to the enzymatic activation of PARP1/2 (35). Notably, PARP1 in a complex with HPF1 specifically promotes ADP-ribosylation at Ser residues of PARP1, histone proteins, and other chromatin-associated factors (34, 36, 37). Whether PARP1-HPF1-dependent ADP-ribosylation is involved in different types of DNA repair or DNA replication is presently unclear.

Here, using cell-free *Xenopus* egg extracts, we have shown how the LIG3-XRCC1 complex functions as a backup DNA ligase in the absence of LIG1. Defective Okazaki fragment ligation results in the accumulation of nicked DNA on replicating chromosomes and subsequently induces ADP-ribosylation of chromatin proteins, including histone H3. ADP-ribosylation upon LIG1 inactivation occurs in a manner dependent on both PARP1 and HPF1. We also demonstrate that inhibition of PARP1 and HPF1 compromises alternative Okazaki fragment ligation by LIG3-XRCC1.

## MATERIAL AND METHODS

Each experiment was performed at least twice. All oligonucleotide sequences are listed in Supplementary Table 1.

### *Xenopus* interphase egg extracts and purification of chromatin

*Xenopus* laevis was purchased from Kato-S Kagaku and handled according to the animal care regulations at the University of Tokyo. Interphase egg extracts were prepared as described previously (38). Unfertilized *Xenopus* eggs were dejellied in 2.5% thioglycolic acid-NaOH (pH 8.2), and were washed three times in 0.2× MMR buffer [5 mM HEPES-KOH (pH 7.6), 0.1M NaCl, 2 mM KCl, 0.1 mM EDTA, 1 mM MgCl_2_, 2 mM CaCl_2_]. After activation in 1× MMR supplemented with 0.3 μg/ml calcium ionophore, eggs were washed 4 times with EB buffer (10 mM HEPES-KOH pH 7.7, 100 mM KCl, 0.1 mM CaCl_2_, 1 mM MgCl_2_, 50 mM sucrose). Packed eggs were crushed by centrifugation (BECKMAN, Avanti J-E, JS13.1 swinging rotor) for 20 min at 18973 × g. Egg extracts were supplemented with 50 μg/ml cycloheximide, 20 μg/ml cytochalasin B, 1 mM DTT, 2 μg/ml aprotinin and 50 μg/ml leupeptin and clarified for 20 min at 48400 × g (Hitachi, CP100NX, P55ST2 swinging rotor). The cytoplasmic extracts were aliquoted and stored at −80°C. Chromatin purification after incubation in egg extracts was performed as previously described with modifications (39). Sperm nuclei were incubated in egg extracts supplemented with an ATP regeneration system (20 mM phosphocreatine, 4 mM ATP, 5 μg/ml creatine phosphokinase) at 3000-4000 nuclei/μL at 22°C. Aliquots (15 μL) were diluted with 150-200 μL chromatin purification buffer (CPB; 50 mM KCl, 5 mM MgCl_2_, 20 mM HEPES-KOH, pH 7.7) containing 0.1% NP-40, 2% sucrose and 2 mM NEM. After incubating on ice for 5 min, extracts were layered over 1.5 ml of CPB containing 30% sucrose and centrifuged at 15,000 × g for 10 minutes at 4°C. Chromatin pellets were resuspended in 1× Laemmli sample buffer, heated for 5 min and analyzed by SDS-PAGE. To inhibit DNA replication, aphidicolin was added to egg extracts at a final concentration of 30 μM.

### Antibodies and immunodepletions

Rabbit polyclonal antibodies raised against *Xenopus* LIG1 were raised against a bacterially expressed recombinant protein encoding a His10-tagged 420-amino-acid fragment from the N-terminus of *Xenopus* LIG1. FEN1 antibodies were raised against the recombinant His10-tagged full-length protein. LIG3 antibodies were raised against a bacterially expressed recombinant protein encoding a C-terminal fragment containing amino acids 638-903 of xLIG3. XRCC1 antibodies were raised against an N-terminus fragment of *Xenopus* XRCC1 encoding amino acids 1-313, which was tagged on the N terminus with His10. xHPF1 and xARH3 antibodies were raised against a bacterially expressed full-length protein which was tagged on the N terminus with His10. PARP1 antibodies used for immunodepletion were raised against a fragment containing amino acids 500-650 of xPARP1. PARG antibodies used for Immunoblotting were raised against a bacterially expressed N-terminus fragment containing 1-385 of xPARG which was tagged on the N terminus with His10. The protein fragments were purified from bacteria grown by Hokudo under native conditions. Histone H3 antibodies were obtained from Abcam (ab1791). Antibodies against PCNA were obtained from SantaCruz (PC10). Pan-ADP-ribose detecting reagent was obtained from Millipore (MABE1016). Antibodies against RPA34 were a generous gift of M. Mechali (Institute of Human Genetics, CNRS). For xPARP1 or xHPF1 depletion, 250 μl of antiserum were coupled to 50 μl of recombinant protein A-sepharose (rPAS, GE Healthcare). Antibody beads were washed three times in PBS and treated with 5 μl fresh rPAS. Beads were washed twice in CPB, split into three portions, and 100 μl extracts were depleted in three rounds at 4°C, each for 1 hour. For xLIG1 and xXRCC1 depletion, 170 μl of antiserum were coupled to 35 μl of rPAS. Antibody beads were washed three times in PBS and treated with 4 μl fresh rPAS. Beads were washed twice in CPB, split into two portions, and 100 μl extracts were depleted in two rounds at 4°C, each for 1 hour.

### Measurement of DNA replication efficiency in *Xenopus* egg extracts

α-^32^P dCTP (3,000 Ci/mmol) was added to an interphase extract containing sperm nuclei, and incubated at 22°C. At each time point, extracts were diluted in reaction stop solution (1% SDS, 40 mM EDTA) supplemented with Proteinase K (NACALAI TESQUE, INC.), followed by overnight incubation at 37°C. The reaction solution was spotted on a glass filter, and then precipitated with a 5% TCA solution containing 2% pyrophosphate. Filters were washed twice with 5% TCA, and then twice with ethanol. After filters were dried, incorporation of radioactivity was counted in scintillation liquid. Radiolabeled dCTP in chromosomal DNA was also monitored by autoradiography following alkaline agarose gel electrophoresis.

### Immunoprecipitation from *Xenopus* egg extracts

5 μl of antiserum were diluted with 250 μL of PBS containing 0.05% NP-40 and coupled to 10 μl of Protein A-Agarose beads (Roche) by rotating at 4°C overnight. Egg extracts were diluted fivefold with CPB buffer, and cleared by centrifugation for 10 min at 14, 000 x g. 50-100 μL of the supernatant was incubated with antibody beads for 1-2 hours at 4°C. Beads were then washed 3 times with CPB buffer containing 0.1% Triton X-100. The bead-bound proteins were eluted with SDS sample buffer and analyzed by immunoblotting. In this case, Protein A-HRP was used as the secondary antibody.

### Histone H3 precipitation from chromatin

Immunoprecipitation of histone H3 from the chromatin fraction was performed as described previously (40). Briefly, mock- or LIG1-depleted extracts containing sperm nuclei were diluted fivefold with CPB supplemented with 0.1% NP-40, 2 mM NEM, and 2 mM tannic acid. After incubating on ice for 5 min, extracts were layered over a sucrose step gradient composed of 500 μl CPB buffer containing 30% sucrose and 10 μl 2M sucrose. Chromatin fractions were resuspended in a 50 μl sucrose cushion, and then supplemented with 50 μl of 2 × nuclease solution [40 mM K-HEPES (pH 7.5), 200 mM KCl, 5 mM MgCl_2_], 20% Triton-100, 20 mM CaCl_2_, MNase (200 U/ml)] and incubated at 22°C for 20 min. The reaction was stopped by the addition of EDTA to 10 mM (final concentration), and the supernatant was obtained by centrifugation at 15,000 rpm for 10 minutes and used as a chromatin-solubilized sample. SDS was added to the supernatant to a final concentration of 1 %, and diluted with tenfold Lysis buffer (1% Triton X-100, 1 mM EDTA, 150 mM NaCl, 15 mM Tris-HCl, pH 8.0). The chromatin lysates were incubated for 2 hours at 4°C with 2 μg of anti-histone H3 antibody conjugated to 10 μl of Protein A-agarose beads. After the beads were washed with lysis buffer, the bound protein was eluted with SDS sample buffer and analyzed by immunoblotting.

### Protein expression and purification

His10-xLIG1 (1-420), His10-xLIG3 (683-903), His10-xXRCC1 (1-313), His10-xPARP1 (600-750), His10-xPARG (1-385), His10-xARH3 (full-length), and His10-xHPF1 (full-length) were expressed and purified from bacteria cells. Escherichia coli BL21-CodonPlus-RIL strain (Agilent) was transformed with a pET19b vector inserted with a DNA fragment encoding a target protein sequence. Cells were cultured in LB medium until the OD600 reached 0.6. The culture was supplemented with 0.25 mM IPTG for 16 hours at 20°C. Cells were harvested and washed twice with ice-cold PBS, and then lysed in 10 mL lysis buffer (50 mM NaH_2_PO_4_, 300 mM NaCl, 10 mM imidazole, pH 8.0) supplemented with 1% NP-40, 2 μg/ml aprotinin, 5 μg/ml leupeptin, 100 μg/ml PMSF, 20 μg/ml trypsin inhibitor by sonication and cleared by centrifugation at 5,000 rpm for 10 min. Cleared lysates were incubated with Ni-NTA agarose (QIAGEN) at 4°C for 2 hours. The agarose beads were washed with lysis buffer containing 20 mM imidazole. Bead-bound proteins were eluted with lysis buffer containing 500 mM imidazole. His10-HPF1 was further applied to a PD-10 desalting column (GE Healthcare BioSciences) equilibrated with EB containing 1 mM DTT.

xLIG1-3×Flag was expressed and purified from insect cells. Briefly, *Xenopus* LIG1 with a C-terminal FLAG x 3 tag was cloned into pVL1392 using primers No. 3 and No.4. F8A/F9A and K721A mutations in pKS104-xLIG1 constructs were introduced using a KOD-Plus Mutagenesis kit (Toyobo).

Baculoviruses were produced using a BD BaculoGold Transfection kit and a BestBac Transfection kit (BD Biosciences), following the manufacturer’s protocol. Proteins were expressed in Sf9 insect cells by infection with viruses expressing wild type xLIG-3×Flag, or its mutant for 72 hours. Sf9 cells were lysed by resuspension in lysis buffer (20 mM Tris-HCl, pH 8.0, 100 mM KCl, 5 mM MgCl_2_, 10% glycerol, 1% Nonidet P40 (NP-40), 1 mM DTT, 10 μg/ml leupeptin, and 10 μg/ml aprotinin). A soluble fraction was obtained after centrifugation of the lysate at 15,000×g for 15 min at 4°C. The fraction was incubated for 4 hours at 4°C with 250 μl of anti-FLAG M2 affinity resin (Sigma-Aldrich) equilibrated with lysis buffer. The beads were collected and washed with 10 ml wash buffer (20 mM Tris-HCl, pH 8.0, 100 mM KCl, 5 mM MgCl_2_, 10% glycerol, 0.1% NP-40, 1 mM DTT) and then with 5 ml EB (20 mM HEPES-KOH, pH 7.5, 100 mM KCl, 5 mM MgCl_2_) containing 1 mM DTT. The recombinant xLIG1 was eluted twice in 250 μl EB containing 1 mM DTT and 250 μg/ml 3×FLAG peptide (Sigma-Aldrich).

Eluates were pooled and concentrated using a Vivaspin 500 (GE Healthcare Biosciences, VS0121). 3×Flag-xLIG3/xXRCC1-myc was purified from insect cells as described above. pVL1392-3×Flag-xLIG3 and pVL1392-xXRCC1-myc were co-transfected and expressed in Sf9 insect cells for 72 hours. *In vitro* translation was performed using a TNT SP6 Quick Coupled Transcription / Translation System (Promega, L2080). 40 μl of rabbit reticulocyte lysates supplemented with methionine were incubated with 1 μg of pKS103-xLIG1, pKS104-xLIG1, pKS103-xLIG3, or pKS103-xXRCC1 for 90 min at 30°C. Lysates expressing recombinant protein were frozen with liquid nitrogen and stored at −80°C.

### Okazaki fragment analysis

Genomic DNA was isolated from egg extracts using a Wizard Genomic DNA purification kit (Promega, A1120). 0.5 μg of DNA were used in labeling reactions in end-label buffer (0.5 mM dNTPs, 50 mM Tris-HCl pH 8.0, 10 mM MgCl_2_) and 10 U Klenow (exo-) polymerase (NEB, M0212S) and α-^32^P-dCTP (Perkin Elmer) at 37°C for 1 hour. The reaction mixture was treated with an equal volume of 2x alkaline loading buffer (100 mM NaOH, 2 mM EDTA, 2.5% Ficoll, 0.025% bromocresol green). Labelled DNA was separated in 1.2% denaturing agarose gels (50 mM NaOH, 1 mM EDTA) and run at 50 V and 30 mA for 7 hours. In order to prevent the loss of small fragments (<150 bp), the gel was fixed by incubation with 7% TCA for 30 min on a shaker, dried and then exposed to phosphor screens or X-ray film.

## RESULTS

### LIG1 is a ligase responsible for Okazaki fragment joining in *Xenopus* egg extracts

We used a cell-free system derived from *Xenopus* egg extracts to delineate mechanisms of Okazaki fragment joining in vertebrates. We first examined whether LIG1 is required for ligation of Okazaki fragments in egg extracts. We cloned the full-length cDNA of *Xenopus* LIG1 (xLIG1) encoding a 1,070 amino acid protein with significant homology to human LIG1 (hLIG1) (88% amino acid similarity) (Supplementary Figure 1A). A fragment of the xLIG1 encoding amino acids 1-420 was purified as a 10 tandem His-tag fused protein and used for specific antigen preparation. The antiserum specifically recognized an approximately 160 kDa band in extracts as well as a full-length xLIG1 recombinant protein generated by in vitro translation with 3×Flag-tag (3×Flag-xLIG1 or xLIG1-3×Flag) (Supplementary Figure 1B).

Consistent with previous reports using budding yeast and mammalian cells, our antibodies co-immunoprecipitated xLIG1 with xPCNA from extracts (Figure 1A). A pull-down experiment using a recombinant wild-type xLIG1-3× Flag showed a tight interaction between xLIG1 and xPCNA whereas that using its mutant within a PIP-box did not (Figure 1B and C). In addition, we found that xLIG1 as well as xPCNA and xFEN1 accumulated on chromatin during S phase. This binding was completely abolished by the addition of a DNA polymerase inhibitor, aphidicolin (APH) (Figure 1D and Supplementary Figure 1C). These results indicate that xLIG1 accumulates on S-phase chromatin in a DNA replication-dependent manner.

**Figure 1.**
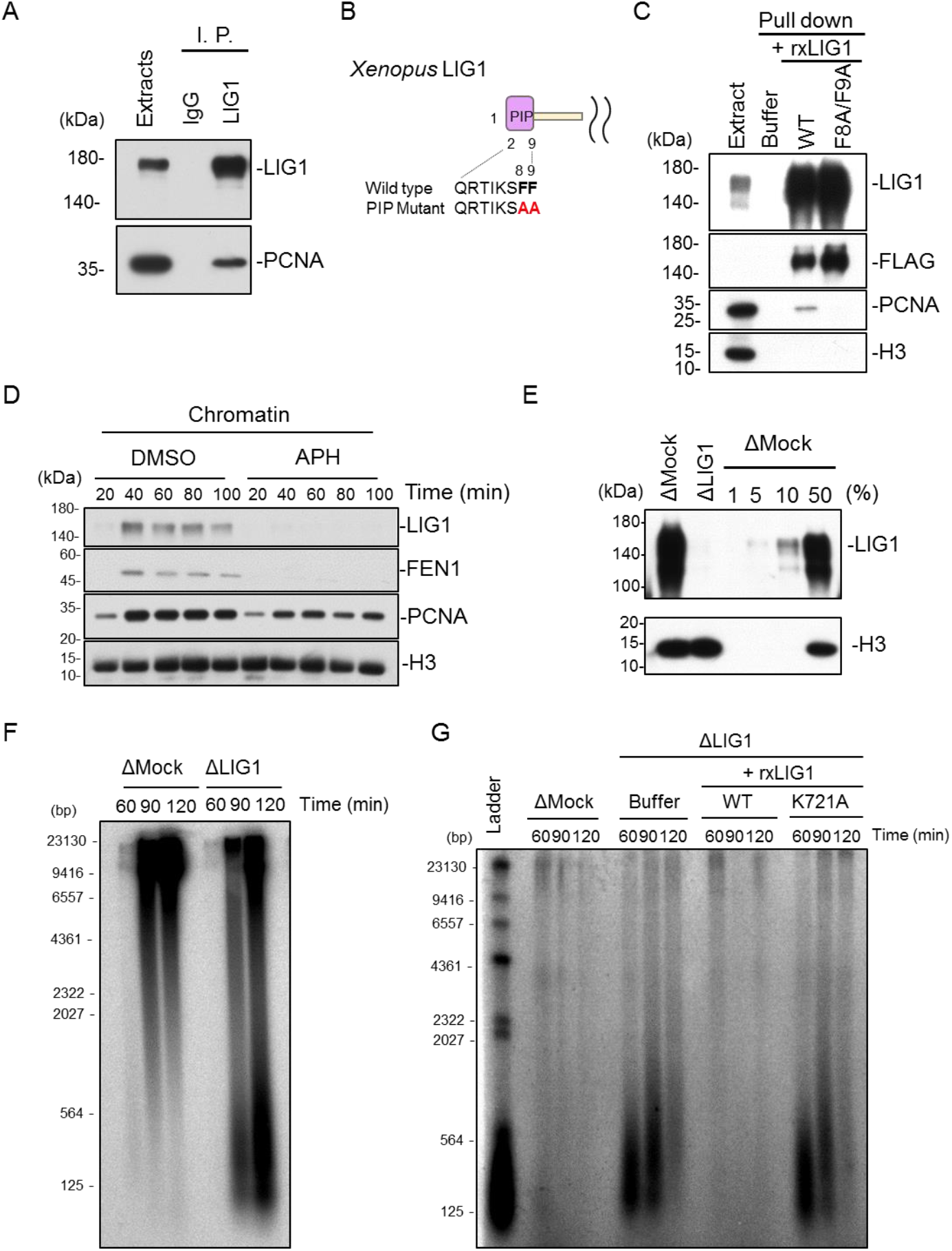
Depletion of LIG1 inhibits Okazaki fragment ligation in *Xenopus* egg extracts. (A) Control or xLIG1 antibodies were used for immunoprecipitation (I. P.) and immunoprecipitated proteins were analyzed by immunoblotting using the indicated antibodies. (B) Sequences of the wild-type and the F8A/F9A mutant PIP-box in xLIG1 are shown. (C) Recombinant wild-type xLIG1-3×Flag or xLIG1-F8A/F9A-3×Flag was added to the egg extracts. Immunoprecipitation using anti-Flag antibodies was performed and resultant immunoprecipitates were analyzed by immunoblotting using the indicated antibodies. (D) Interphase egg extracts were treated with sperm chromatin and incubated in the presence of 30 μM aphidicolin or in its absence (DMSO). Chromatin fractions were isolated and analyzed by immunoblotting using the indicated antibodies. (E) Immunodepletion efficiency of xLIG1 from *Xenopus* egg extract. (F) Replication products in mock- or xLIG1-depleted extracts were separated by alkaline agarose gel electrophoresis followed by autoradiography. (G) xLIG1-depleted extracts were supplemented with wild-type xLIG1-3×Flag or xLIG1-K721A-3×Flag and chromatin was isolated. Purified genomic DNA from chromatin was labeled using exonuclease-deficient Klenow fragment and α-^32^P dCTP, and separated in a denaturing agarose gel.

To test whether LIG1 is required for Okazaki fragment joining in *Xenopus* egg extracts, we immunodepleted xLIG1 from egg extracts and examined the accumulation of small DNA fragments during S phase which was initiated by the addition of sperm DNA. More than 99% of xLIG1 was depleted without a significant effect on gross DNA replication of sperm chromatin (Figure 1E and Supplementary Figure 1D). However, denaturing agarose gel analysis of DNA replication products revealed the accumulation of short DNA fragments in xLIG1-depleted extracts, but not in mock-depleted extracts, presumably a result of a failure of Okazaki fragment ligation (Figure 1F). We then labeled unligated Okazaki fragments using DNA polymerase and α-^32^P-dCTP. In mock-depleted extracts, nicked DNA was almost undetectable. In contrast, xLIG1 depletion resulted in a transient, but marked accumulation of short nascent DNA fragments (Figure 1G). The defective joining of Okazaki fragments was rescued by the addition of wild-type xLIG1 recombinant protein (Figure 1G). In contrast, a mutant xLIG1 lacking enzymatic activity failed to join Okazaki fragments (Figure 1G) although the chromatin binding of the mutant was comparable to that of the wild-type (Supplementary Figure 1E and F). Reintroduction of recombinant mutant xLIG1 (R794L and R924W), a point mutant that was identified in an immunodeficient patient (41, 42), also showed a defect in Okazaki fragment joining similar to that of the inactive mutant, and retained chromatin binding activity (Supplementary Figure 1G-K). A previous study in yeast showed that completion of Okazaki fragment joining triggers ELG1/ATAD5-RFC-dependent PCNA unloading from chromatin (43). Consistent with this, immunodepletion of xLIG1 resulted in increased chromatin binding of xPCNA, and xFEN1 (Supplementary Figure 1L and M). These results confirm the importance of the DNA ligase activity of LIG1 in Okazaki fragment joining in *Xenopus* egg extracts.

### A LIG3-XRCC1 complex acts to compensate for Okazaki fragment ligation for LIG1 function

Notably, the accumulation of nicked DNA in LIG1-depleted extracts was diminished with the progression of S phase (Figure 1G), suggesting the existence of a compensatory pathway other than that of LIG1 for Okazaki fragment joining in egg extracts. Previously, the viability of cells lacking LIG1 was reported to depend on LIG3 function (15, 16). Therefore, we examined the possibility that the LIG3-XRCC1 complex acts as part of a compensatory system for the Okazaki fragment joining. A comparison of nucLIG3-α and XRCC1 between humans and *Xenopus* revealed conserved domain structures with 61-88% and 70-78% homology, respectively (Supplementary Figure. 2A). To clarify the function of *Xenopus* LIG3-XRCC1 complex, we raised antibodies against xLIG3 and xXRCC1. These antibodies recognized polypeptides of the expected molecular weight in egg extracts as well as their recombinant proteins expressed in reticulocyte lysates as a control (Supplementary Figure. 2B). We showed that xLIG3 and xXRCC1 stably interact in *Xenopus* egg extracts and that xXRCC1 immunodepletion resulted in an efficient co-depletion of xLIG3 (Figure. 2A, Supplementary Figure 2B). We then investigated whether xLIG3 contributes to Okazaki fragment joining during S-phase. Depletion of xXRCC1 or xLIG1/xXRCC1 did not have any significant effect on gross DNA replication (Supplementary Figure 2C). As mentioned above, the accumulation of Okazaki fragments was observed in xLIG1-depleted extracts, but not in xLIG3-xXRCC1-depleted extracts. In contrast, when xLIG1 and xLIG3-xXRCC1 were simultaneously depleted, the amount of nicked Okazaki fragments was markedly increased and was not eliminated over S phase (Figure 2B). We then examined the chromatin binding of xLIG3-xXRCC1 during DNA replication. Both xLIG3 and xXRCC1 were detected on chromatin in mock-depleted extracts. When xLIG1 was depleted, a marked increase in chromatin binding of xLIG3 and xXRCC1 was detected, particularly during late S phase (Figure 2C, Supplementary Figure 2D). In contrast, xXRCC1 depletion had little effect on the recruitment of xLIG1 to chromatin. Reintroduction of recombinant xLIG3-xXRCC1 complex purified from insect cells revealed a chromatin loading of xLIG3 and xXRCC1 similar to that of endogenous proteins, and rescued impaired Okazaki fragment joining (Figure 2D and E, Supplementary Figure 2F). Substitution of arginine and lysine in conserved PAR-binding sites of xXRCC1 with alanine (Supplementary Figure 2E) resulted in inefficient Okazaki fragment joining as compared to wild type-xXRCC1, although it had no clear effect on xLIG3-xXRCC1 chromatin binding (Figure 2D and E). Together, our results suggest that LIG3-XRCC1 is responsible for Okazaki fragment joining when LIG1 function is perturbed.

**Figure 2.**
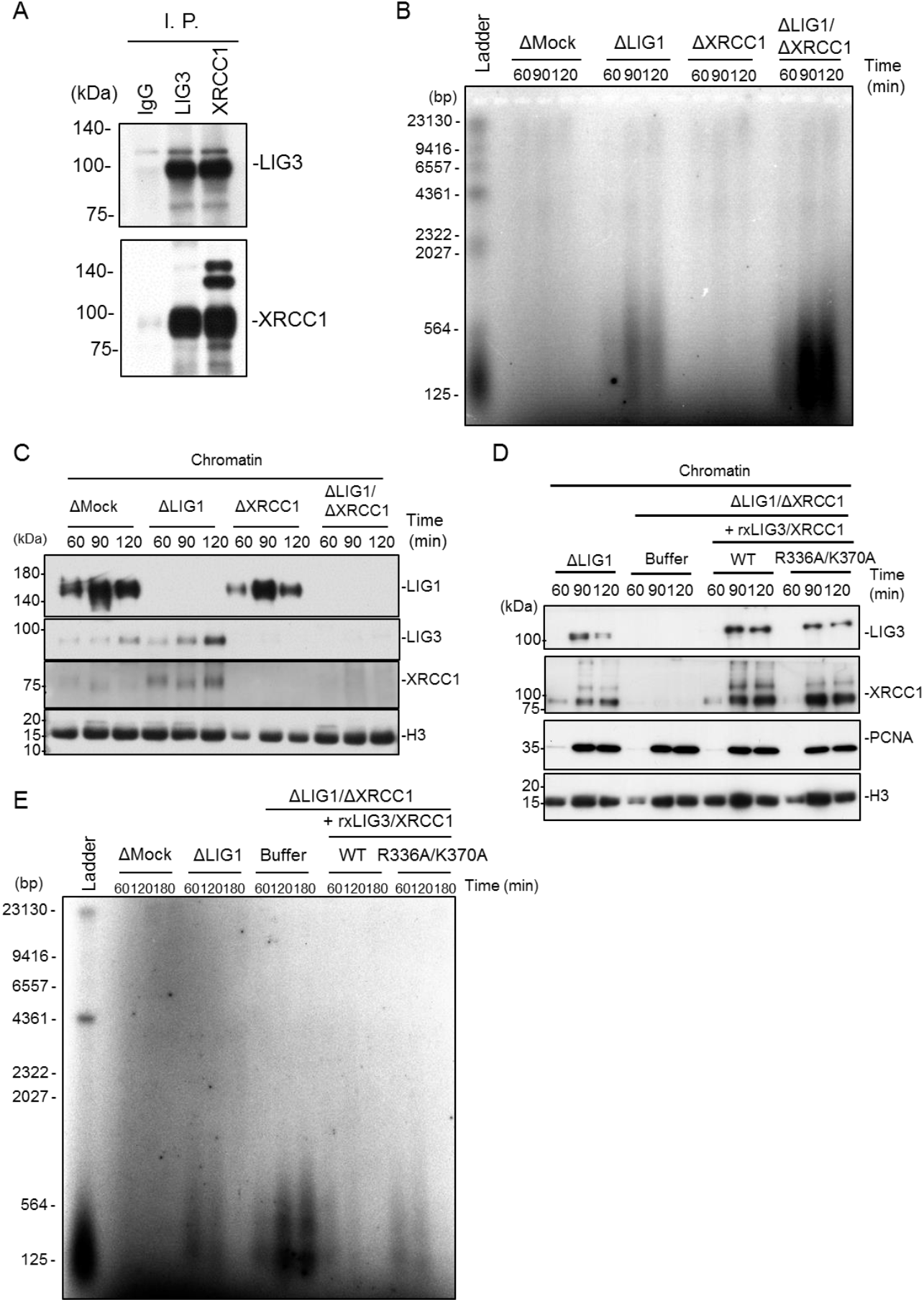
LIG3-XRCC1 ensures Okazaki fragment ligation in the absence of LIG1. (A) Immunoprecipitation (I. P.) of xLIG3 and xXRCC1 from egg extracts. I. P. was performed with anti-xLIG3 or -xXRCC1 antiserum. Pre-immune serum was also used as a control. Immunoprecipitates were analyzed by immunoblotting using the indicated antibodies. (B) Mock-, xLIG1-, xXRCC1-, or xLIG1/xXRCC1-depleted extracts were used to replicate sperm chromatin. Genomic DNA was purified and analyzed as in Figure 1G. (C) Chromatin-bound proteins from (B) were monitored using immunoblotting using the indicated antibodies. (D) xLIG1- and xLIG1/xXRCC1-depleted extracts were used to replicate sperm chromatin. xLIG1/xXRCC1-depleted extracts were supplemented with either buffer (+Buffer), wild-type 3×Flag-xLIG3-xXRCC1-myc, or 3×Flag-xLIG3-xXRCC1-R336AK370A-myc lacking the poly-ADP-ribose binding activity of xXRCC1. (E) Replication products from (D) were analyzed as in Figure 1G.

### HPF1-dependent PARP activation promotes alternative Okazaki fragment ligation

It has been demonstrated that LIG3-XRCC1 is recruited to single-strand break sites in a PARP1-mediated poly-ADP-ribosylation dependent manner (23–25). Recent studies also revealed that ADP-ribosylation at serine residues is a major modification upon DNA damage (34–36). ADP-ribosylation at serine is known to be catalyzed by PARP1 in complex with HPF1, which targets histone proteins and PARP1 per se as a substrate (33–35). We therefore compared the ADP-ribosylation of mock- or xLIG1-depleted chromatins using an anti-pan-ADP-ribose binding reagent, which recognizes both mono- and poly-ADP-ribose. ADP-ribosylation signals were readily detectable in xLIG1-depleted extracts, but not in mock-depleted extracts. Importantly, reintroduction of recombinant wild-type xLIG1, but not its enzymatically inactive mutant, fully suppressed these modifications (Figure 3A, Supplementary Figure 3A). Similar signals were detected when xLIG1 (R794L) and xLIG1 (R924W) mutants were reintroduced in xLIG1-depleted extracts (Supplementary Figure 3B and C). These bands were increased by co-depletion of xPARG with xLIG1, suggesting that they were ADP-ribosylated proteins (Figure 3B). Depletion of xPARG alone from egg extracts had only a slight effect on ADP-ribosylation of chromatins. Thus, ADP-ribosylation is likely to be induced by LIG1 dysfunction. In mammalian cells, apart from Okazaki fragment joining, LIG1 is known to be involved in various DNA repair pathways both during and outside of S phase (7). Importantly, we identified a protein band of 15 kDa as a major target of ADP-ribosylation after xLIG depletion. Since previous reports showed histone H3 to be a molecular target of PARP1-HPF1-dependent ADP-ribosylation (33, 44), we examined whether the 15 kDa band corresponded to histone proteins. Immuoprecipitation of histone H3 revealed an ADP-ribosylation signal in anti-H3 immunoprecipitates from xLIG1-depleted, but not mock-depleted chromatin (Figure 3C). We found that ADP-ribosylation in xLIG1-depleted extracts was completely abolished by the addition of APH (as seen by an increased loading of xRPA34) (Figure 3D, Supplementary Figure 3D). The results suggest that histone H3 ADP-ribosylation upon xLIG1-depletion is dependent on DNA replication rather than not the indirect consequence of the accumulation of spontaneous DNA damage or replication fork stalling by xLIG1 depletion. HPF1-dependent ADP-ribosylation is reported to be hydrolyzed by ADP-ribose hydrolase 3 (ARH3). Thus, ADP-ribosylation in xLIG1-depleted extracts might be sensitive to ARH3. When an excess amount of recombinant wild-type xARH3 from bacteria cells, as well as its mutant lacking enzymatic activity (D85N, corresponding to hARH3 D77N) were added to xLIG1-depleted extracts (0.2~1 μM), only wild-type recombinant ARH3 diminished histone H3 ADP-ribosylation (Figure 3E, Supplementary Figure 3E). These results suggest that LIG1 deficiency leads to activation of PARP1-HPF1-dependent ADP-ribosylation targeting histone H3.

**Figure 3.**
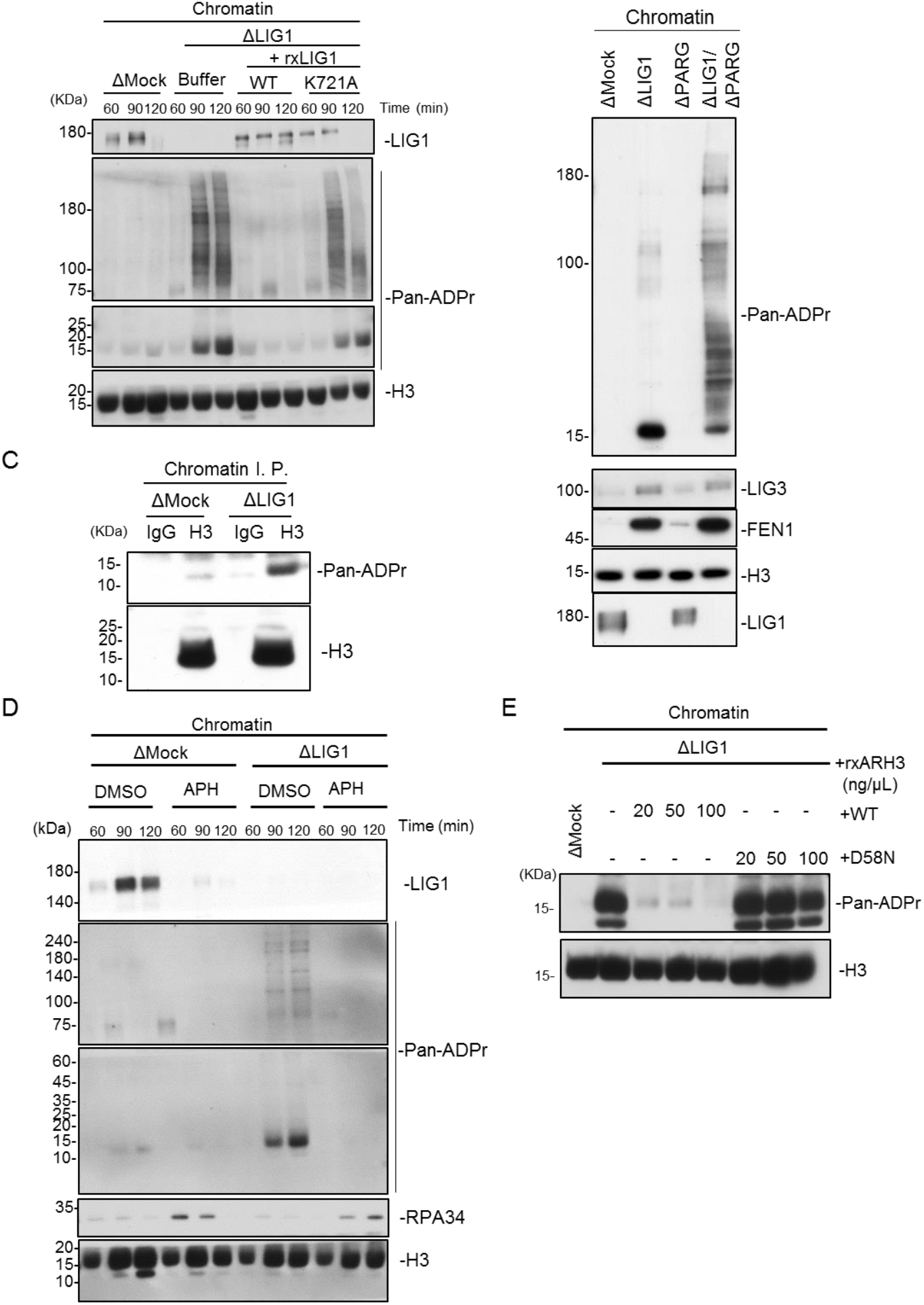
LIG1 deficiency causes histone H3 ADP-ribosylation. (A) Mock- and xLIG1-depleted extracts were used for replication of sperm chromatin. xLIG1-depleted extracts were supplemented with wild-type xLIG1 or xLIG1-K721A. Chromatin-bound proteins were analyzed by immunoblotting using the indicated antibodies, and pan ADP-ribose detecting reagent. (B) Mock-, xLIG1-, xPARG-, and xLIG1/xPARG-depleted extracts were used for replication of sperm nuclei. Chromatin-bound proteins were analyzed as in Figure 3A. (C) Chromatin fractions from the indicated immunodepleted extracts were solubilized with MNase followed by immunoprecipitation with anti-histone H3 antibodies after treatment with 1% SDS. The resultant immunoprecipitates were analyzed as in Figure 3A. (D) Mock- and xLIG1-depleted extracts were used for replication of sperm nuclei in the presence of 30 μM APH where indicated. Chromatin-bound proteins were analyzed by immunoblotting using the indicated antibodies, and pan ADP-ribose detecting reagent. (E) Mock- and xLIG1-depleted extracts were used for replication of sperm chromatin. xLIG1-depleted extracts were supplemented with either buffer or recombinant wild-type His10-xARH3 or His10-xARH3-D58N, (0.2~1 μM). Chromatin-bound proteins were analyzed by immunoblotting using the indicated antibodies, and pan ADP-ribose detecting reagent.

To directly test whether ADP-ribosylation is mediated by PARP1 and HPF1, we performed immunodepletion of PARP1 and HPF1 in the absence of LIG1. Under this condition, immunodepletion of xPARP1 partially inhibited histone H3 ADP-ribosylation on xLIG1-depleted chromatin (Figure 4A and B), perhaps because of compensation by PARP2 or other nuclear PARPs (30). xHPF1 depletion resulted in a more pronounced suppression of histone H3 ADP-ribosylation (Figure 4C and D). This defect was largely restored by recombinant wild-type xHPF1, but not by a xHPF1 mutant harboring mutations in the conserved acidic corner (alanine substitution of Tyr251 and Arg252, corresponding to hHPF1 Try238 and Arg239), which disrupts the interaction between xHPF1 and xPARP1 (Figure 4E and F) (35). These results indicate that the PARP1-HPF1 complex stimulates histone H3 ADP-ribosylation in the absence of LIG1.

**Figure 4.**
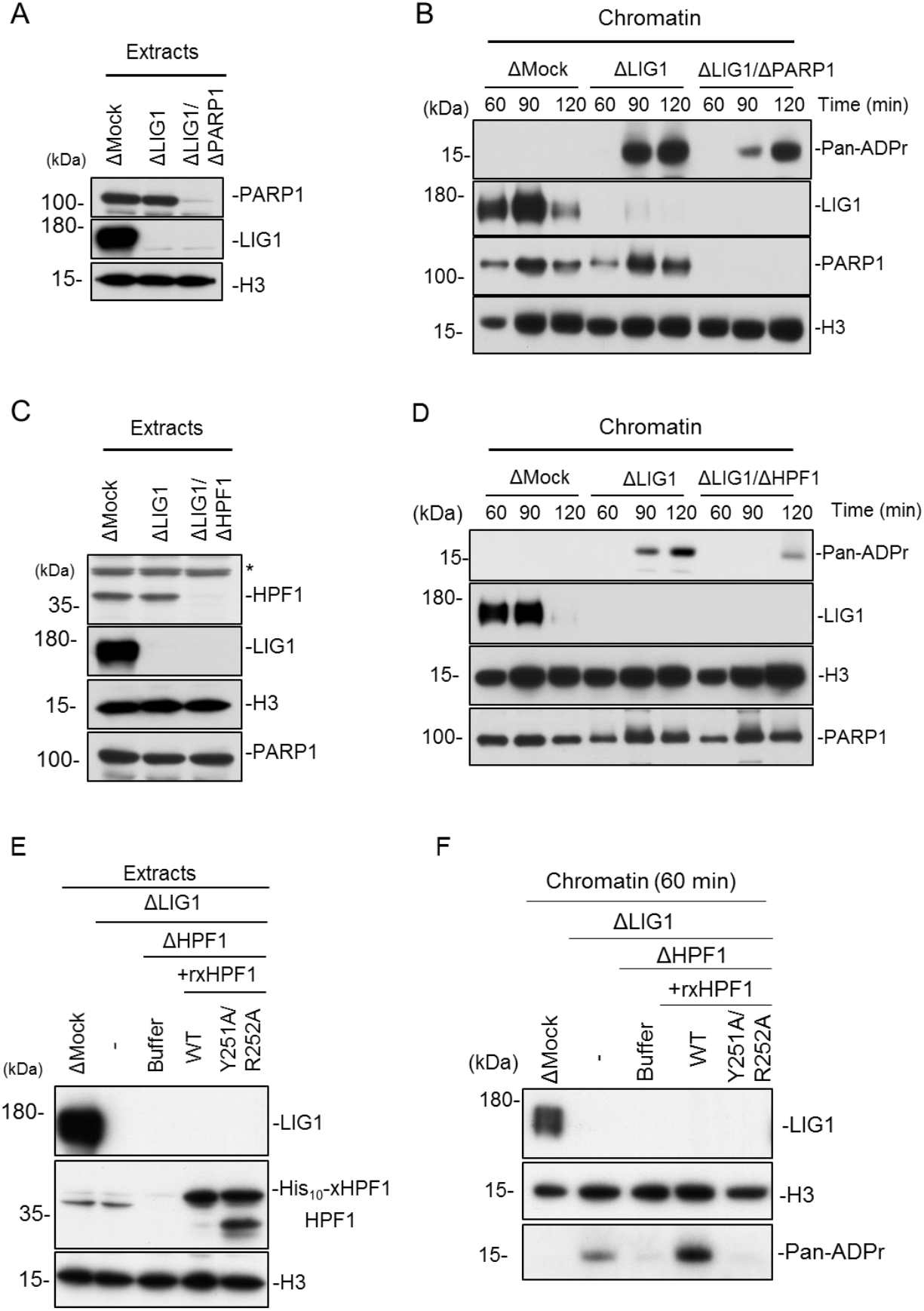
PARP1-HPF1 stimulates histone H3 ADP-ribosylation in LIG1-depleted extracts. (A) Mock-, xLIG1-, and xLIG1/xPARP1-depleted extracts were analyzed by immunoblotting using the indicated antibodies. (B) The extracts from (A) were used to replicate sperm nuclei. Chromatin-bound proteins were analyzed by immunoblotting. (C) Mock-, xLIG1-, and xLIG1/xHPF1-depleted extracts were analyzed by immunoblotting using the indicated antibodies. The asterisk indicates a non-specific band. (D) The extracts from (C) were used to replicate sperm nuclei. Chromatin-bound proteins were analyzed. (E) xLIG1/xHPF1-depleted extracts were supplemented with buffer, recombinant His10-xHPF1 (WT), or recombinant mutant His10-xHPF1 (Y251AR252A) defective for PARP1 binding activity. (F) The extracts from (E) were used to replicate sperm nuclei. Chromatin-bound proteins were analyzed.

Based on the above findings, we hypothesized that HPF1-dependent ADP-ribosylation promotes LIG3-XRCC1-dependent Okazaki fragment ligation. We found that xLIG3 displayed decreased chromatin binding in the absence of xPARP1 which was correlated with a loss of ADP-ribosylation (Figure 5A, Supplementary Figure 4A). Similarly, the depletion of xHPF1 also inhibited the chromatin binding of xLIG3-xXRCC1 (Figure 5B, Supplementary Figure 4B). Consistent with this observation, unligated Okazaki fragments in xLIG1-depleted extracts further increased with xPARP1, or xHPF1 co-depletion (Figure 5C). Collectively, our data suggest that LIG1 deficiency causes PARP1-HPF1-dependent ADP-ribosylation, which acts to activate LIG3-XRCC1 as a backup pathway for Okazaki fragment ligation.

**Figure 5.**
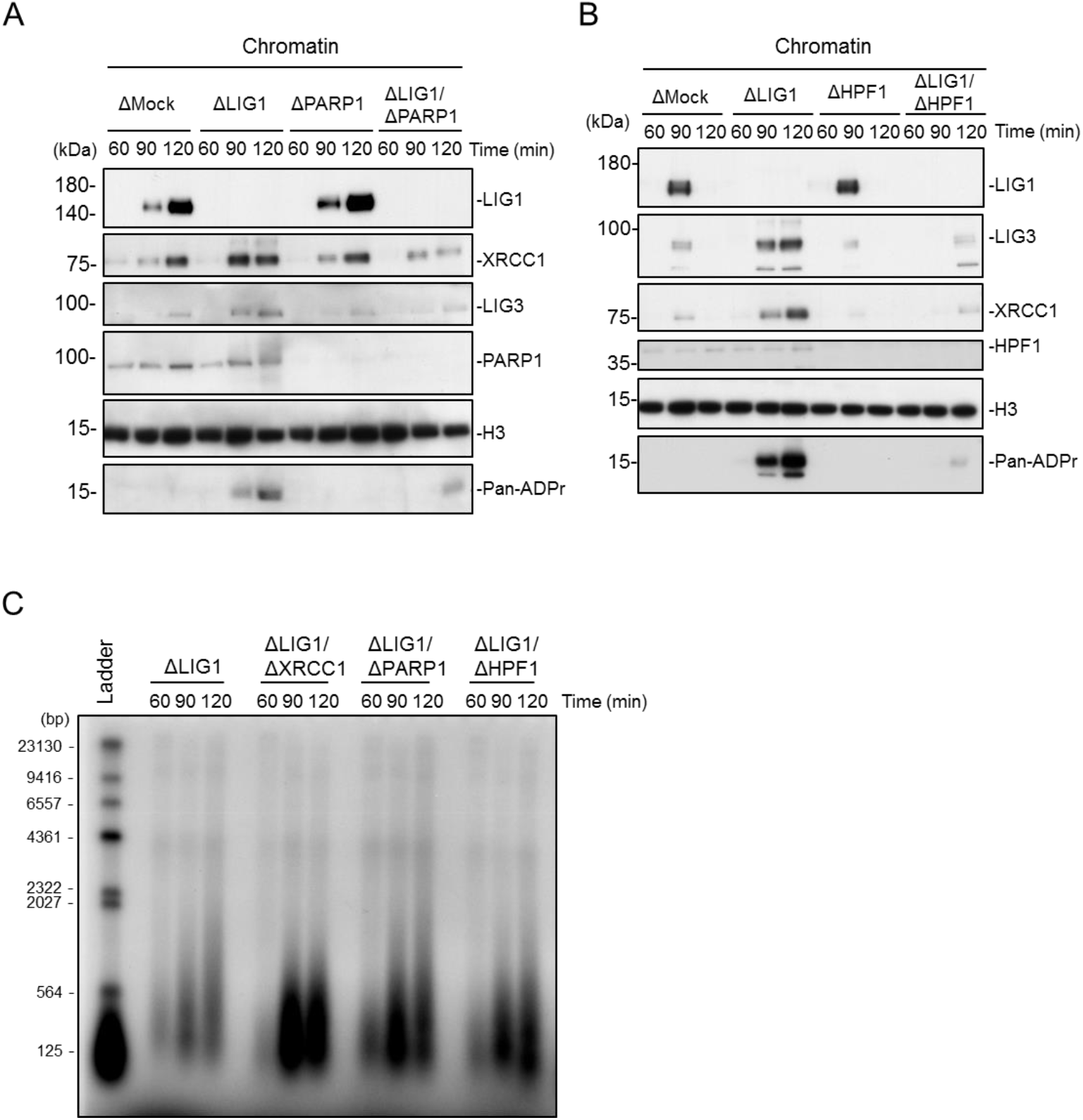
HPF1-dependent PARP activation promotes alternative Okazaki fragment ligation. (A) Mock-, xLIG1-, xPARP1-, and xLIG1/xPARP1-depleted extracts were used for replication of sperm nuclei. Chromatin-bound proteins were analyzed by immunoblotting using indicated antibodies. (B) Mock-, xLIG1-, xHPF1-, and xLIG1/xHPF1-depleted extracts were used for replication of sperm nuclei. Chromatin-bound proteins were analyzed by immunoblotting using the indicated antibodies. (C) LIG1-, xLIG1/xXRCC1-, xLIG1/xPARP1-, or xLIG1/xHPF1-depleted extracts were used to replicate sperm chromatin. Genomic DNA was purified and labeled using exonuclease-deficient Klenow fragments and α-32P dCTP. Samples were separated in a denaturing agarose gel and incorporation of radioactivity was monitored by autoradiography.

## DISCUSSION

Co-depletion of LIG1 and LIG4 did not significantly affect cell proliferation (14–16) whereas that of LIG1 and LIG3 induced lethality, suggesting the redundant function of LIG3 in Okazaki fragment ligation. However, the regulatory mechanisms for how LIG1 and LIG3 function properly during S phase were not clear. Using the *Xenopus* cell-free replication system, we found that unligated Okazaki fragment accumulates in early to mid-S phase, but is eventually resolved in late S phase in the absence of LIG1. LIG3-XRCC1 accumulates on chromatin specifically in late S phase in the LIG1 depleted extracts. Co-depletion of LIG1 and LIG3-XRCC1 resulted in a complete loss of Okazaki fragment joining. Finally, PARP1-HPF1 is essential for LIG3-XRCC1 in Okazaki fragment joining. Consistent with this, LIG1 inhibition was reported to activate PARP at replication sites in mammalian cells (45) although involvement of this PARP activation in LIG3 recruitment has not yet been examined. Previously, XRCC1 was reported to co-localize with PCNA during S phase under unperturbed conditions (21, 22, 45). Taken together, our results suggest that LIG1 might be frequently perturbed during normal DNA replication, resulting in chromatin sites with impaired Okazaki fragment joining where LIG3-XRCC1 has to be recruited. Supporting this idea, recent genome-wide nucleotide resolution mapping of the 3’-OH terminus (GLOE-seq) suggested that the complementary actions of LIG1 and LIG3 control the fidelity of Okazaki fragment ligation in human cells (46). Although the precise mechanism of LIG3-XRCC1 chromatin recruitment remains to be determined, our results and those of a previous report (45) suggest that PARP1-HPF1-dependent ADP ribosylation is important for this event. With respect to the activation of PARP1 in LIG1-depleted extracts, PARP1-HPF1 might directly recognize unligated Okazaki fragments through the DNA-binding domain of PARP1 (31), which in turn activate PARP1.

PARP activity is important for the recruitment of LIG3-XRCC1 during DNA repair (30), and inhibition of LIG1 and FEN1 activates PARP and promotes PAR synthesis at DNA replication sites (45, 47) although the substrate proteins and the mode of ADP ribosylation have not yet been determined. We found that histone H3 is the main substrate for ADP ribosylation by PARP1-HPF1. Consistent with this, the immunodepletion of PARP1 or HPF1 attenuated LIG3-XRCC1 function in Okazaki fragment joining. Although our results suggest a possible role of PARP1-HPF1-dependent histone H3 ADP-ribosylation in the chromatin recruitment of LIG3-XRCC1, it does not exclude their alternative roles. For example, ADP-ribosylation inhibits the conversion of nicked DNA into double-stranded DNA breaks by preventing DNA2 or EXO1-dependent resection (48, 49). ADP-ribosylation is also known to promote ALC1-mediated chromatin remodeling, presumably through the ADP ribosylation of histone H3 (33, 50, 51). Intriguingly, PARP1-HPF1 specifically catalyzes ADP ribosylation at a Ser residue. Therefore, LIG3-XRCC1 as well as other factors might have unidentified domains which specifically recognize histone H3 ADP ribosylation. Synergistic anti-cancer effects of histone deacetylase (HDAC) and PARP inhibitors are reported in a variety of cancers (52, 53). Our current work thus provides an important cue to develop a therapeutic strategy for cancers with FEN1, POL δ, and LIG1 mutations in which Okazaki fragment joining is strongly suppressed upon treatment with PARP and HDAC inhibitors (54).

## SUPPLEMENTARY DATA

Supplementary Data are available at NAR online.

## ACKNOWLEDGEMENT

We thank Mitsuko Masutani (Nagasaki University) for helpful discussions.

