## Supplemental Data for "HPF1-dependent PARP activation promotes LIG3-XRCC1-mediated backup pathway of Okazaki fragment ligation"

**Figure S1**

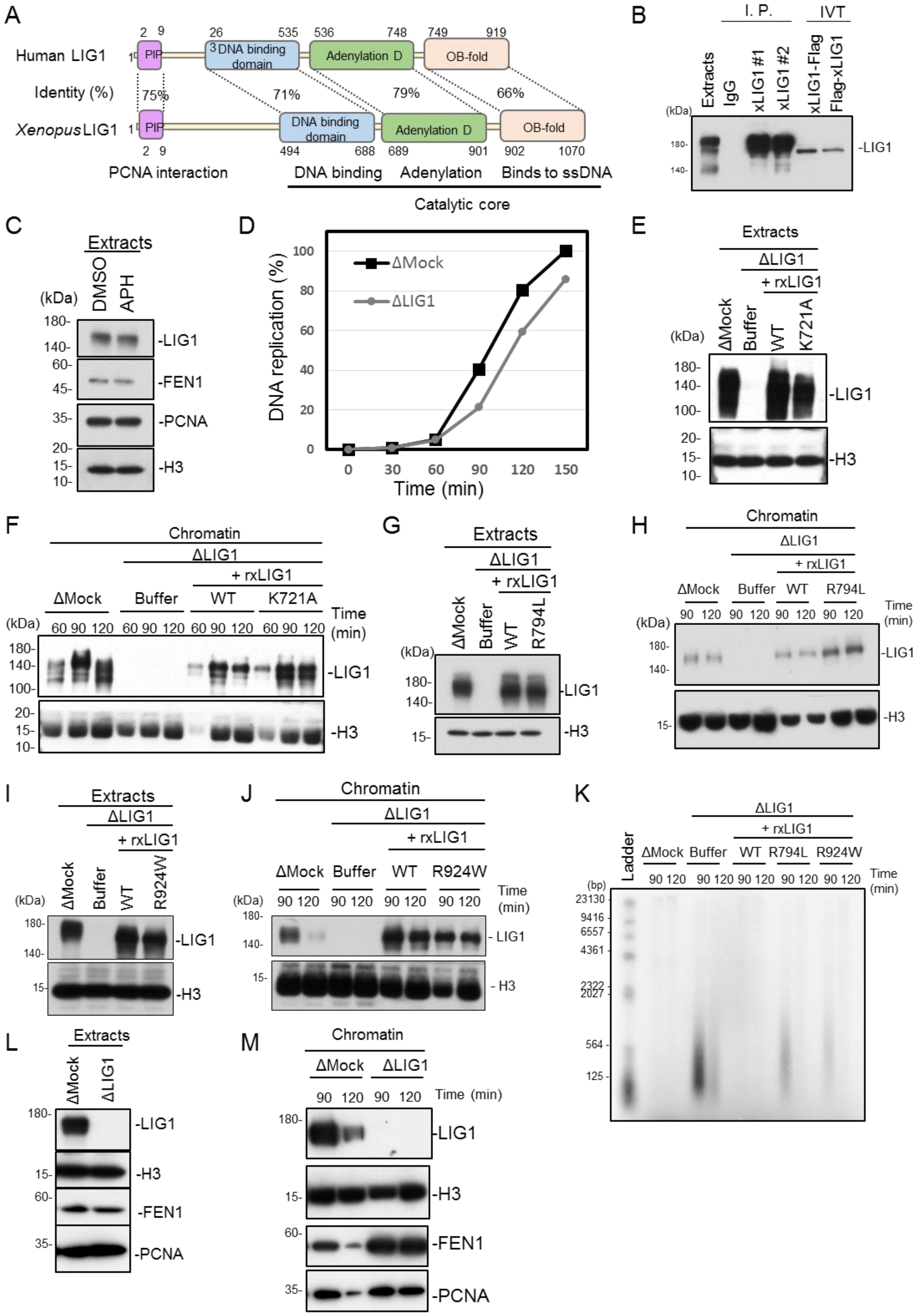
A
Human LIG1
Identity (%)
Xenopus LIG1
PIP
DNA binding domain
Adenylation D
OB-fold
75%
71%
79%
66%
2 9
26 535 536 748 749 919
2 9
494 688 689 901 902 1070
PCNA interaction
DNA binding
Adenylation
Binds to ssDNA
Catalytic core
B
I. P.
IVT
Extracts
IgG
xLIG1 #1
xLIG1 #2
xLIG1-Flag
Flag-xLIG1
(kDa)
180-
140-
-LIG1
C
Extracts
DMSO
APH
-LIG1
-FEN1
-PCNA
-H3
D
ΔMock
ΔLIG1
DNA replication (%)
Time (min)
E
Extracts
ΔLIG1
+ rxLIG1
ΔMock
Buffer
WT
K721A
F
Chromatin
60 90 120
Time (min)
G
R794L
H
I
R924W
J
K
Ladder
(bp)
23130
9416
6557
4361
2322
2027
564
125
L
M

Supplementary Figure 1. (A) Domain structures of *Xenopus* and Human LIG1. LIG1 is composed of a PCNA-interacting protein (PIP) motif, a DNA binding domain, an adenylation domain and an oligonucleotide/oligosaccharide-binding (OB)-fold domain that are well conserved in humans and *Xenopus*. (B) Immunoblot analysis of egg extracts, LIG1 immunoprecipitates and reticulocyte lysates translating recombinant FLAG-tagged xLIG1. (C) Immunoblot of extracts used in Figure 1D using the indicated antibodies. (D) DNA replication in xLIG1- and mock-depleted extracts. The relative amounts of DNA synthesis are shown. (E) LIG1-depleted extracts were supplemented with wild-type xLIG1 or xLIG1-K721A and analyzed by immunoblotting using the indicated antibodies. (F) The extracts from (E) were used to replicate sperm nuclei. Chromatin-bound proteins were analyzed by immunoblotting. (G) LIG1-depleted extracts were supplemented with wild-type xLIG1 or xLIG1-R794L and analyzed by immunoblotting using the indicated antibodies. (H) The extracts from (G) were used to replicate sperm nuclei. Chromatin-bound proteins were analyzed by immunoblotting. (I) LIG1-depleted extracts were supplemented with wild-type xLIG1-3×Flag or xLIG1-K924W-3×Flag and analyzed by immunoblotting using the indicated antibodies. (J) The extracts from (G) were used to replicate sperm nuclei. Chromatin-bound proteins were analyzed by immunoblotting. (K) xLIG1-depleted extracts were supplemented with wild-type xLIG1-3×Flag, xLIG1-R794L-3×Flag or xLIG1-R924W-3×Flag. Purified genomic DNA from chromatin was labeled using exonuclease-deficient Klenow fragment and $\alpha$-$^{32}$P dCTP and separated in a denaturing agarose gel. (L) Mock- and xLIG1-depleted extracts were analyzed by immunoblotting using the indicated antibodies. (M) The extracts from (L) were used to replicate sperm nuclei. Chromatin-bound proteins were analyzed by immunoblotting.

**Figure S2**

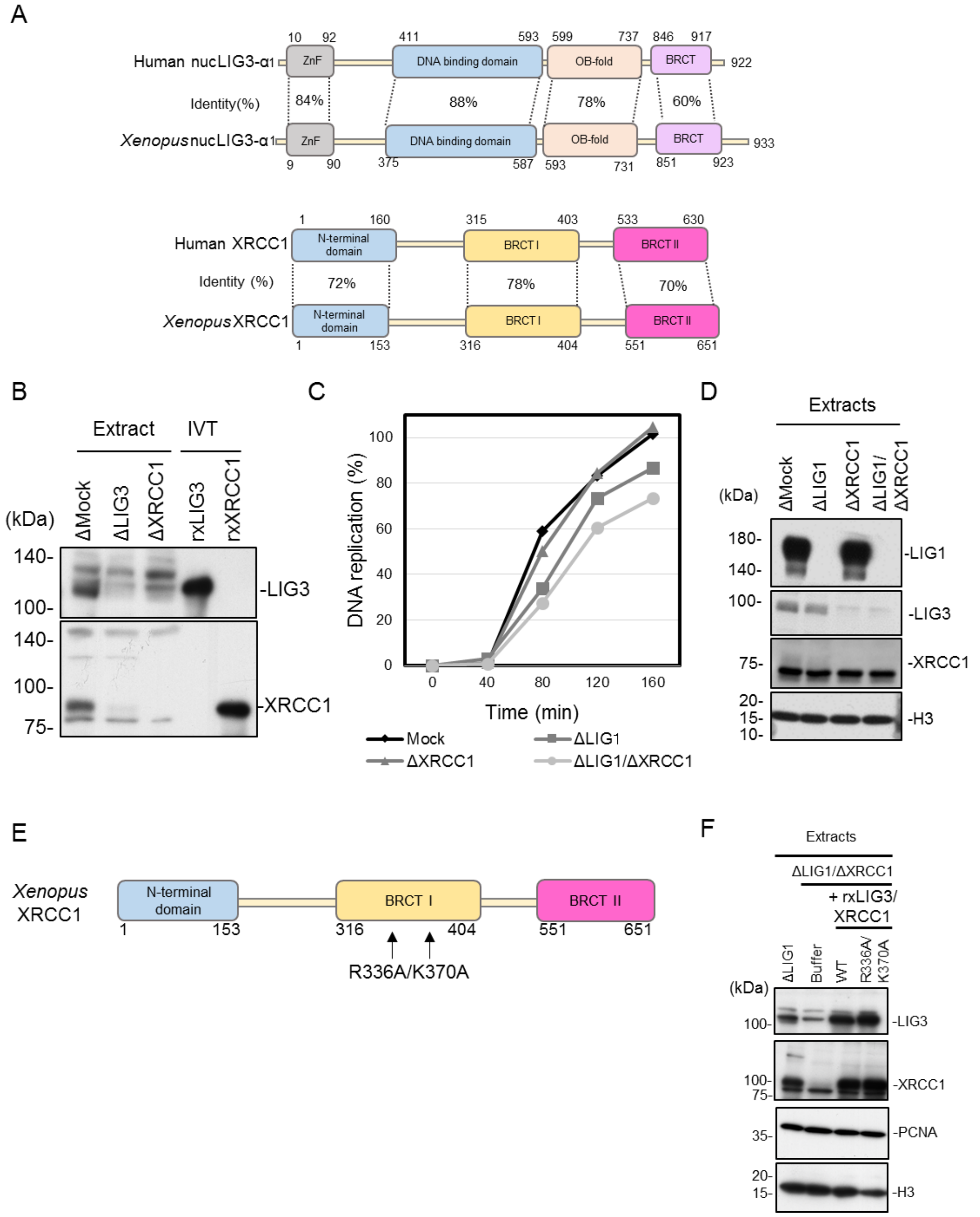


Supplementary Figure 2. (A) Domain structures of *Xenopus* and Human nuclear LIG3 and XRCC1. Nuclear LIG3 is composed of a zinc-finger (ZnF), DNA binding, oligonucleotide/oligosaccharide-binding (OB)-fold and BRCA1 C-Terminal (BRCT) domains that are well conserved in humans and *Xenopus*. XRCC1 is composed of an N-terminal domain, and BRCT I and BRCT II domains that are well conserved in humans and *Xenopus*. (B) Immunodepletion efficiency of xLIG3 and xXRCC1 from *Xenopus* egg extract. IVT; in vitro translation protein. Depleted extracts and IVT proteins were analyzed by immunoblotting using the indicated antibodies. (C) DNA replication in xLIG1-, xXRCC1-, xLIG1/xXRCC1-, and mock-depleted extracts. The relative amounts of DNA synthesis are shown. (D) Immunoblot of the extracts of Figure 2C using the indicated antibodies. (E) Sequences of the wild-type and R336A/K370A mutant BRCT I domain in xXRCC1 are shown. (F) Immunoblot of the extracts of Figure 2D using the indicated antibodies.

**Figure S3**

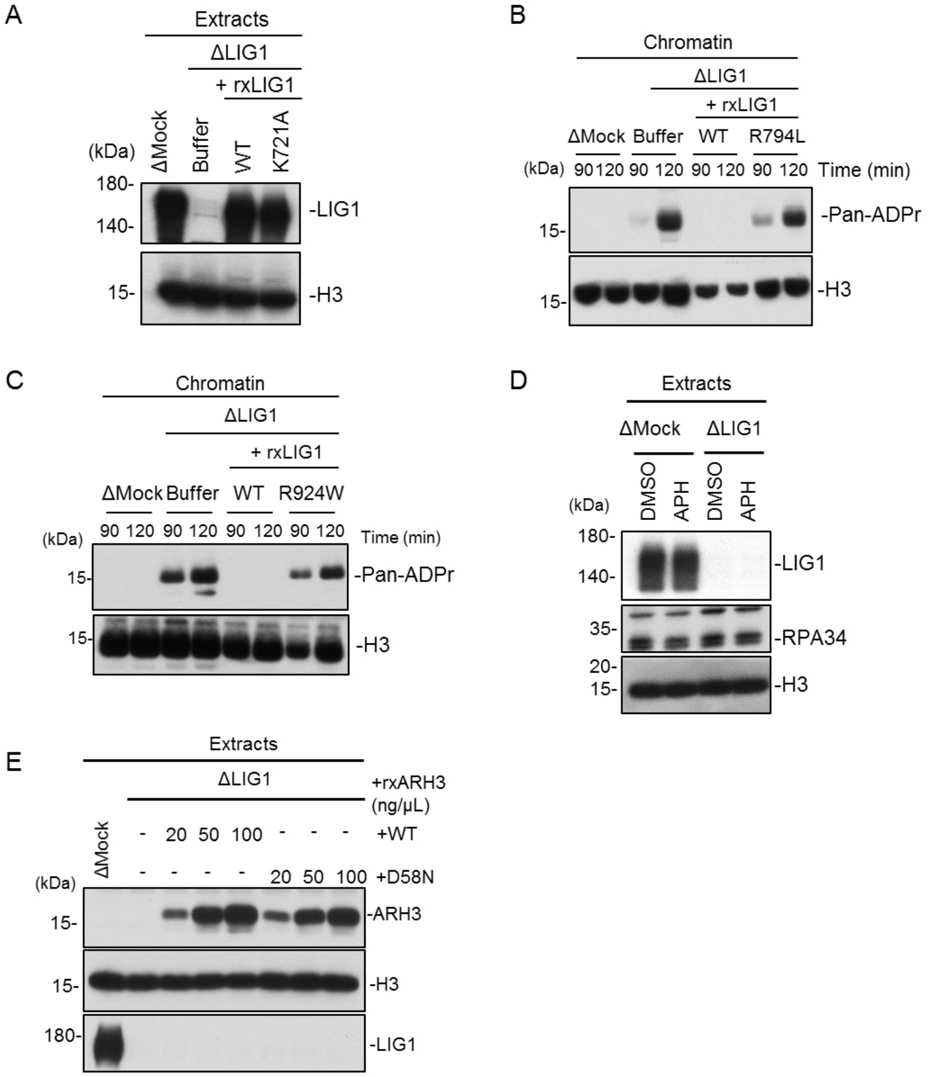


Supplementary Figure 3. (A) Immunoblot of extracts used in Figure 3A using the indicated antibodies. (B) xLIG1-depleted extracts were supplemented with either buffer or wild-type xLIG1-3×Flag or xLIG1-3×Flag-R794L. Chromatin-bound proteins were analyzed by immunoblotting using pan ADP-ribose detecting reagent. (C) xLIG1-depleted extracts were supplemented with either buffer or wild-type xLIG1-3×Flag or xLIG1-3×Flag-R924W. Chromatin-bound proteins were analyzed by immunoblotting using pan ADP-ribose detecting reagent. (D) Immunoblot of the extracts of Figure 3D using the indicated antibodies. (E) Immunoblot of the extracts of Figure 3E using the indicated antibodies..

**Figure S4**

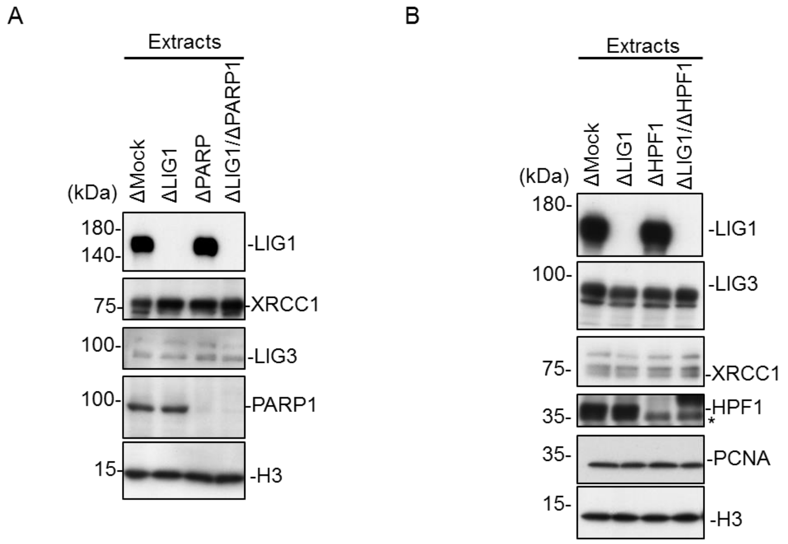


Supplementary Figure 4. (A) Immunoblot of the extracts of Figure 5A using the indicated antibodies. (B) Immunoblot of the extracts of Figure 5B using the indicated antibodies. The asterisk indicates a non-specific band.

**Table S1**

Supplementary Table 1

| No. | Sequence 5'-3' | Description |
|---|---|---|
| 1 | GAAACCTTCATTTTACGGCGGGGA | xLIG1 amplification |
| 2 | ACTCCAAGGCAACACAATAGGTGG | xLIG1 amplification |
| 3 | GGCGCGGATCAGATCTCATGCAACGAACAATAAAGTC | pVL139-xLIG1-Flag3 |
| 4 | GGGCCCTCTAGAATTCTACTTGTTATCGTCATCCT | pVL139-xLIG1-Flag3 |
| 5 | GCATATGACGGGGAGCGTGCACAGATAC | xLig1K721A mutation |
| 6 | GTATTCACAAGTAAAGGCAGCTTCA | xLig1K721A mutation |
| 7 | GCTGCTCAGCCCAAGCTAGGGGCTGAAGTAA | xLig1F8AF9A mutation |
| 8 | GGACTTTATTGTTCGTTGCATGAGA | xLig1F8AF9A mutation |
| 9 | CTAAAGAGAAAGGATGTGGATGCATCAG | xLig1R794L mutation |
| 10 | AGTAGTCAGTACTTGAAATGGCTGA | xLig1K794L mutation |
| 11 | TGGACTGGTATCTATGGAGGCTTCTTAC | xLig1R924W mutation |
| 12 | TTTCCCTTTCCCAAGGTAAGCTCCA | xLig1R924W mutation |
| 13 | GAGTGGGCTTTCTCTGTGTTGTTG | xFEN1 amplification |
| 14 | GACATATTGTCAAAGCCTTAGCGC | xFEN1 amplification |
| 15 | TGTGTGAGTGAGGGAAATTGCAGG | xLIG3 amplification |
| 16 | GTTGATCTGTGATGCCTTCTCCTG | xLIG3 amplification |
| 17 | GGGCCCTCTAGAATTTTACACACAATTCTCATCAACAC | pVL139-xLIG3-Flag3 |
| 18 | GGTGGTGTGAAGCAGAAATCTGCT | xXRCC1 amplification |
| 19 | CTTTGCACTGGCAACATCCAGTTC | xXRCC1 amplification |
| 20 | GGCGCGGATCAGATCTCATGCCTGTGATCAAACTGAAGC | pVL139-xXRCC1-Flag3 |
| 21 | GGGCCCTCTAGAATTCGCCTTGGGCACCACAACGTAAGG | pVL139-xXRCC1-Flag3 |
| 22 | GCAGCCGACCTCCGTGATAAAGCATTAG | xXRCC1R336A mutation |
| 23 | GAATGGATTCTGAAAGCCGCTCAGC | xXRCC1R336A mutation |
| 24 | GCATTCAGCCAGGTGAAAGCAGCGGGCG | xXRCC1R370A mutation |
| 25 | AGGGGTATTTGCAAAGGCACATATG | xXRCC1R370A mutation |
| 26 | CACGTTGTTACTGTAAGAGCCAGTGTGTTCCCTGT | xPARP1 amplification |
| 27 | CACTCCGTGATCCATCCTGCCCAACGTATG | xPARP1 amplification |
| 28 | AGCGGAAACTCCACTCTTGGTTGA | xPARG amplification |
| 29 | GCTGGACATGAGTCAGGACTTCAT | xPARG amplification |
| 30 | TTCTCAGAGAGTGCTGCTATTCGC | xHPF1 amplification |
| 31 | AAGCCATCTTCCCTCTGAAGTGAC | xHPF1 amplification |
| 32 | GCTGCGGAGCTTCCAGAGACAGATGGAAACC | xHPF1Y251AR252 mutation |
| 33 | CCCAACATCATTCTTATCCACTGGC | xHPF1Y251AR252 mutation |
| 34 | TTCTCAGAGAGTGCTGCTATTCGC | xARH3 amplification |
| 35 | AAGCCATCTTCCCTCTGAAGTGAC | xARH3 amplification |
| 36 | TGTATATGTTAAAACACTCTTTAGT | xARH3D58N mutation |
| 37 | AATGACACAGCCATGGCAAGGTCGATTG | xARH3D58N mutation |
